# A draft reference genome sequence for *Scutellaria baicalensis* Georgi

**DOI:** 10.1101/398032

**Authors:** Qing Zhao, Jun Yang, Jie Liu, Meng-Ying Cui, Yuming Fang, Wengqing Qiu, Huiwen Shang, Zhicheng Xu, Yukun Wei, Lei Yang, Yonghong Hu, Xiao-Ya Chen, Cathie Martin

## Abstract

*Scutellaria baicalensis* Georgi is an important medicinal plant used worldwide. Information about the genome of this species is important for scientists studying the metabolic pathways that synthesise the bioactive compounds in this plant. Here, we report a draft reference genome sequence for *S. baicalensis* obtained by a combination of Illumina and PacBio sequencing, which was assembled using 10 X Genomics and Hi-C technologies. We assembled 386.63 Mb of the 408.14 Mb genome, amounting to about 94.73% of the total genome size, and the sequences were anchored onto 9 pseudochromosomes with a super-N50 of 33.2 Mb. The reference genome sequence of *S. baicalensis* offers an important foundation for understanding the biosynthetic pathways for bioactive compounds in this medicinal plant and for its improvement through molecular breeding.

## Introduction

*Scutellaria baicalensis* Georgi or Chinese Skullcap is a well-known medicinal plant that originated in East Asia and is cultivated in many European countries for its therapeutic properties (Shang et al., 2010). The dried root of *S. baicalensis* has been used as a traditional medicine for more than 2000 years in China, where it has the name Huang-Qin 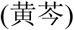 (Zhao et al., 2016). Huang-Qin is used in traditional Chinese medicine (TCM) for treatment of bitter, cold, liver and lung problems as recorded by *Shennong Bencaojing* (*The Divine Farmer’s Materia Medica*), an ancient book on medicine and agriculture, written between about 200 and 250 AD (Ma, 2013). Recent scientific studies have reported the pharmacological activities of preparations of Scutellaria, particularly those of flavonoids that accumulate at high levels in the roots and rhizomes of *S. baicalensis* (Li-Weber, 2009; Shang et al., 2010; Tu et al., 2016). More than 132 compounds have been found in Huang-Qin, including flavones, flavanones, phenylethanoids and their glycosides (Qiao et al., 2016). The flavonoids in roots of Scutellaria have been reported to have various beneficial bioactivities including antibacterial, antiviral, antioxidant, anti-cancer, hepatoprotective and neuroprotective properties (Gao et al., 2011; Yang et al., 2012). Preparations of Scutellaria roots or their extracts are used as ingredients in many drugs for cold and liver problems and as adjunct treatments for lung and liver cancers in both Asian and Western countries (Wen, 2007). Despite the commercial importance and increasing demand for *Scutellaria*, improvements through breeding have been almost non-existent. The absence of genome information for this historically important medicinal plant has limited the understanding of how its flavonoid bioactives are made and prevented the improvement of productivity through genetic selection. Understanding the genes responsible for biosynthesis of the various flavonoids made in *S. baicalensis* and their regulation, will lay a foundation for biosynthesis and molecular breeding for improved productivity

## Results

The DNA for genome sequencing of *Scutellaria baicalensis* Georgi came from a single plant maintained in Shanghai Chenshan Botanical Garden. DNA was extracted and sequenced by Illumina and PacBio sequencing strategy. We obtained 48.02 Gb PacBio reads, amounting to 117.66 X coverage of the 408.14 Mb, a genome size estimated by k-mer distribution analysis (Supplementary Figure 1a and supplementary table 1). We also measured the genome size to be 392 Mb by flow cytometry, which is close to the value given by k-mer method (Supplementary Figure 1b). After interactive error-correction among the PacBio reads, assembly was carried out using FALCON to obtain primary contigs. To avoid problems of heterozygosity from outcrossing, the contigs were phased using FALCON-Unzip, then the updated primary contigs and haplotigs were polished with Quiver. The final contigs were error-corrected with the 67.96 Gb (166.51X) of short reads obtained from Illumina HiSeq X Ten sequencing (Supplementary table 1). The consensus sequences were further assembled using the reference of 86.72 G (212.485 X coverage) from 10 X genomics sequencing (Supplementary table 1). All the contigs were extended using FragScaff to generate an assembly with a total scaffold length of 386.6 Mb (94.73% of the genome) and N50 of 1.33 Mb (Supplementary table 2). To facilitate genome annotation and to obtain expression profiles of the *S. baicalensis* genes, we performed transcriptome sequencing from RNA samples extracted from flowers, flower buds, leaves, stems, roots, and JA-treated roots. RNA samples from each tissue were sequenced in triplicate using the HiSeq 2000 platform.

A Hi-C (in vivo fixation of chromosomes) library was then employed to refine the first version of the reference genome. This method sorted 475 of the 578 scaffolds into 114 super-scaffolds, accounting for 98.04% of the original 386.63 Mb assembly (Table 1 and Supplementary table 3). All the super-scaffolds could be located on 9 groups (Supplementary figure 2). The super-scaffold N50 reached 33.2 Mb, with the longest one being 87.96 Mb (Table 1 and Supplementary table 4). The groups are hereafter referred to as pseudochromosomes, and this number corresponds well to the number of chromosomes reported in previous studies (1n=9, 2n=18) (Cheng et al., 2010). The genome of *S. baicalensis* has a normal A/T/C/G content, and its GC content is 34.24%, with N comprising 0.6% (Supplementary table 5). SNP calling based on the genome sequence shows a heterozygosity rate of 0.31% (Supplementary table 6).

**Table 1.** Overview of *S. baicalensis* draft reference genome assembly.

| Assembly feature | Statistic |
| --- | --- |
| Estimated genome size (k-mer analysis) | 408.14 Mb |
| Assembly length | 386.63 Mb |
| Length loaded on pseudochromosomes | 379.07 Mb |
| Number of pseudochromosomes | 9 |
| Number of super-scaffolds | 114 |
| N50 of super-scaffolds | 33.2 Mb |
| Longest super-scaffolds | 87.96 Mb |
| Assembly % of genome | 94.73 |
| Repeat region of % assembly | 53.95 |
| Predicted gene models | 28 930 |
| Average coding sequence length | 1 122.48 bp |
| Average exons per gene | 5.07 |

To test the coverage of the genome, we mapped the short reads generated from Illumina sequencing, and 96.5% of these reads could be mapped to the scaffolds with a 99.78% overall coverage (Supplementary table 7). EST sequences generated from transcriptome sequencing were also mapped to the assembly and we found that 86.12% ESTs had more than 90% of their sequences in one scaffold, 97.51% ESTs had more than 50% of their sequences in one scaffold (Supplementary table 8). CEGMA and BUSCO evaluations of the genome sequence indicated 96.37% and 94% completeness, respectively (Supplemtal table 9 and Supplental table 10). All data suggest a good quality assembly.

A pipeline combined with *de novo*, homolog and RNA-seq was used to construct gene models for the *S. baicalensis* genome. A total of 28,930 genes were generated this way, with an average length of 2,980 bp and an average CDS length of 1,122 bp (Table 1 and Supplementary table 11), among which, 23,027 genes (79.6%) were supported by RNA-sequencing data. A total of 20,234 genes (69.9%) were supported by all three methods (RNA-seq, de novo and homolog), therefore these genes have been annotated with high-confidence. The resulting protein models were then compared to protein sequences in four protein databases, namely NR, Swiss-Prot, KEGG and InterPro. We found that 28524 (98.6%) gene products could be annotated by at least one of the databases. All the gene loci were named according to the nomenclature used for Arabidopsis, which indicated clearly the relative positions of genes on the pseudochromosomes.

The assembled *S. baicalensis* reference genome contains 55.15% repetitive sequences. Tandem duplications (small satellites and micro-satellites) and interspersed repeats account for 1.2% and 53.95% of the genome, respectively. Long terminal repeats of retroelements (LTR) are the most abundant interspersed repeat, occupying 34.4% of the genome, followed by DNA transposable elements at 15.4% (Supplementary table 12). Genes encoding non-coding RNAs; 1218 miRNAs, 517 tRNAs, 1846 rRNAs and 512 snRNAs were annotated in the genome (Supplementary table 13). An overview of the genes, repeats, non-coding RNA densities and all detected gene duplications are shown in Fig. 1.

**Fig. 1.**
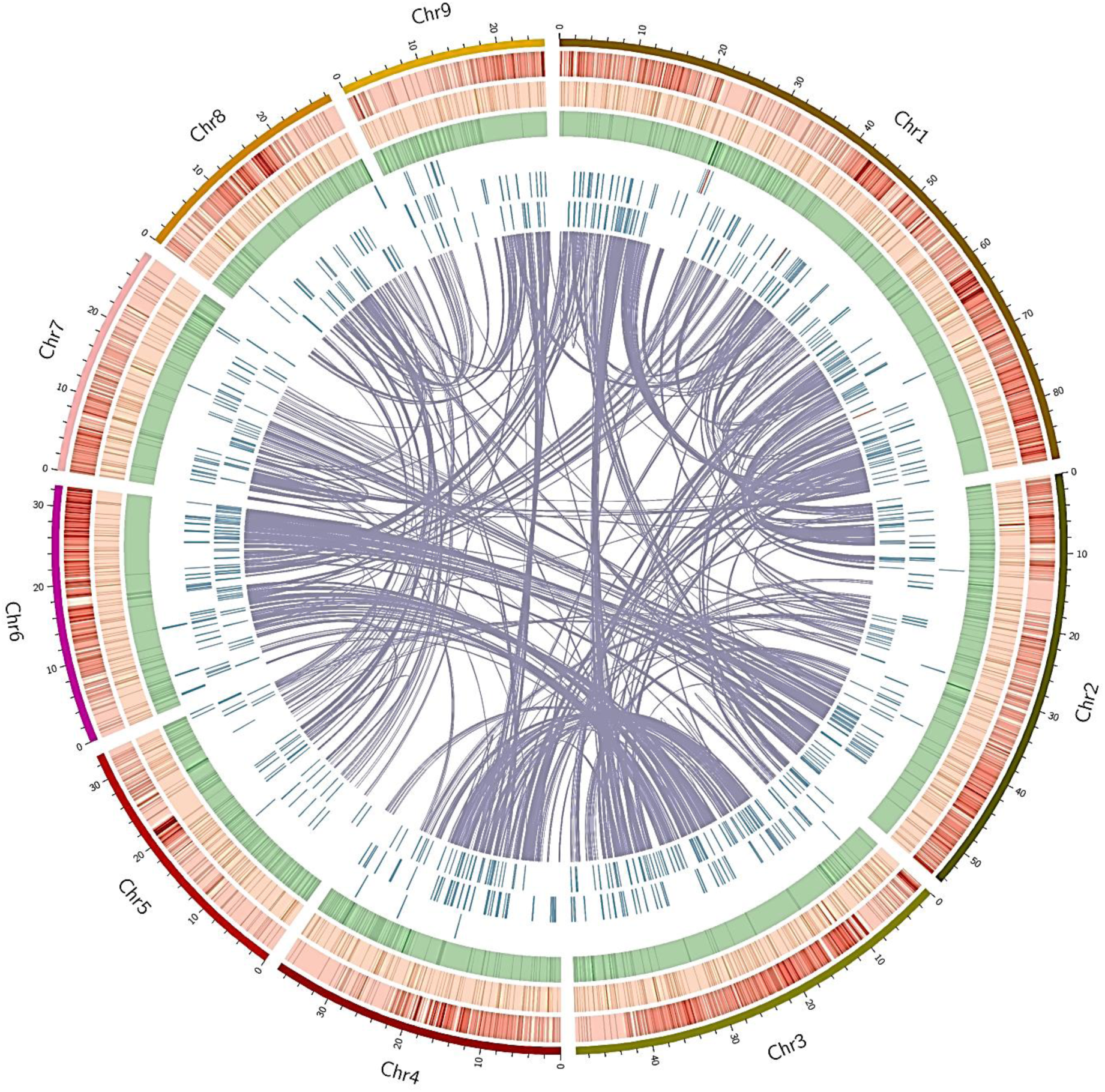
Summary of current *S. baicalensis* draft genome assembly. The outer lines represent pseudochromosomes. The coloured bands summarise the density of genes (red), genes encoding miRNAs (orange), repeats (green), genes encoding rRNAs (blue), genes encoding snRNAs (blue) and genes encoding tRNAs (blue). All detected gene duplications are indicated with links inside the circles.

## Discussion

Members of the family Lamiaceae are commonly known as mint plants. The family has a worldwide distribution, and includes about 6,900 to 7,200 species with the largest genera being Salvia (900 species), followed by Scutellaria (360 species) and Stachys (300 species) (Raymond M. Harley, 2004). The extremely high degree diversity in specialised metabolites makes this family famous as medicinal plants and herbs, including mint, rosemary, basil, savory, perilla, salvia and skullcaps (Scutellaria) and so on. Terpenoids, phenolic acids and flavonoids are mainly responsible for these bioactivities (Lei et al., 2016; Wu et al., 2016; Zhao et al., 2016). Genome sequencing provides a key resource to study the molecular basis of diversity of the specialised metabolites in members of the family Lamiaceae and how their biosynthetic pathways evolved. To date, only the genomes of *S. miltiorrhiza* and *S. splendens* from the family Lamiaceae have been reported (Xu et al., 2016; Dong et al., 2018). We report here a high quality draft reference genome sequence of *S. baicalensis* of 386.63 Mb (about 94%). The genomic information reported offers a foundation for comparative genomic analysis between members of the family Lamiaceae. Furthermore, the genome sequence makes gene isolation and characterization, elucidation of metabolic pathways, as well as molecular breeding, easier for this important medicinal plant

## Methods

### Genome Sequencing

Genomic DNA was extracted from leaves of a single *Scutellaria baicalensis* plant maintained in Shanghai Chenshan Botanical Garden, using a modified CTAB method (Tel-Zur et al., 1999). Quality control was done using a Sage Science Pippin pulse electrophoresis system. Genomic DNA with a length of around 150 kb was sheared using a Megaruptor DNA system and the resulting fragments of 30-50 kb were collected for the following steps. A SMRTbell DNA library was constructed with Sequel 2.0 reagent according to the manufacturer’s instructions (Pacific Bioscience, www.pacb.com). SMRTbell sequencing primers together with P4 DNA polymerase were used for sequencing reactions on SMART Cells. The genomic DNA was sequenced on the PacBio Sequel system. This work produced 48.02 GB of single molecule data, about 117.66 X of the genome (Supplemental Table 1).

For short-insert library construction, DNA was extracted from the same plant using a DNA secure Plant Kit (TIANGEN, http://www.tiangen.com/) according to the manufacturer’s instructions. The DNA was sheared and fragments with sizes of 200-300 bp were retrieved from agarose gels. The fragments were ligated to adaptors and were selected for RNA amplification for templates. The library was then sequenced on HiSeq X Ten. The Illumina sequencing produced 67.96 GB short reads data, amounting to about 166.51 X of the genome.

For 10 X genomic sequencing, we extracted DNA samples using the modified CTAB method (Tel-Zur et al., 1999). The GemCode Instrument (10 X Genomics) was employed for DNA indexing and barcoding. GEM reactions were carried out based on about 1 ng, 50 kb single DNA molecules and 16-bp barcodes were ligated to the molecules in droplets. The intermediate DNA was extracted from the droplets and sheared into 500 bp fragments (Zheng et al., 2016). The fragments were ligated to P7 adaptors, which were then sequenced on Illumina HiSeq X Ten platform (Mostovoy et al., 2016). We obtained 86.72 GB data from 10 X genomics sequencing, which was then used for genome assembly.

DNA from young leaves, collected from *Scutellaria baicalensis*, was used as starting material for the Hi-C library. Formaldehyde was used for fixing chromatin. The leaf cells were lysed and DpnII endonuclease was used for digesting the fixed chromatin. The 5’ overhangs of the DNA were recovered with biotin-labelled nucleotides and the resulting blunt ends were ligated with each other using DNA ligase. Proteins were removed with protease to release the DNA molecules from the cross-links. The purified DNA was sheared into 350 bp fragments and ligated with adaptors (Yaffe and Tanay, 2011). The fragments labelled with biotin were extracted using streptavidin beads and, after PCR enrichment, the libraries were sequenced on Illumina Hiseq PE150.

### Estimation of genome size

The genome size was measured by flow cytometry according to the protocol described by Dolezel (Doležel and BartoŠ, 2005). Briefly, young leaves of *S. baicalensis* were chopped with a sharp razor blade for 60 seconds in nuclear isolation buffer (200 mM TRIS; 4 mM MgCl2.6H2O; 0.5 % (v/v) Triton X-100; pH 7.5). Samples were incubated for 3 minutes, filtered through 50 µm CellTrics^R^ filters. Plant cell nuclei were stained by adding 2 ml buffer containing propidium iodide and RNAse A in the dark for 2 minutes. The relative Nuclear Genome Size was analysed on a flow cytometer (BD FACSAria III). This analysis gave us an estimate of genome size of 392 Mb compared with the tomato genome (Consortium, 2012). We further evaluated the genome size by k-mer frequency analysis based on Illumina short reads using the K-mer Analysis Toolkit (http://www.earlham.ac.uk/kat-tools).

### Genome assembly

De novo assembly was carried out based on PacBio reads. Error correction was conducted by mapping the seed reads in FALCON according to the manufacturer’s instructions (https://github.com/PacificBiosciences/FALCON/wiki/Manual) with the following parameters: -- max_diff 100; --max_cov 100; --min_cov 2; --min_len 5000. Contaminations were removed with PacBio’s whitelisting pipeline. The resulting primary assembly was phased using FALCON-Unzip (default parameters). Heterozygosity of the contigs was analysed with FALCON-Unzip, which was then phased according to the differences. The phased sequences were assembled into haplotigs for a diploid. The contigs were polished with Quiver (http://pbsmrtpipe.readthedocs.io/en/master/getting_started.html) with the parameters: pbsmrtpipe.options.chunk_mode: True pbsmrtpipe.options.max_nchunks: 50, to produce primary contigs, which were further corrected with reference to the Illumina reads with Pilon (https://github.com/broadinstitute/pilon/wiki).

### RNA sequencing and analysis

Six tissues were harvested from 3-month-old *Scutellaria baicalensis* plants, namely, flower buds, flowers, leaves, roots, JA-treated roots (100 µM MeJA treated for 24 hours), and stems. Each tissue was collected for 3 biological replicates. Total RNA was extracted from these tissues using the RNAprep pure Plant Kit (Tiangen). After removing DNA, mRNA was isolated using oligodT beads. The mRNA was harvested and broken into short fragments, which were then used as templates for cDNA synthesis. After end repair, a single nucleotide A (adenine) was added and adapters were ligated to the cDNA. Fragments of 200-300 bp were separated for PCR amplification. An Agilent 2100 Bioanaylzer and ABI StepOnePlus Real-Time PCR System were used for quality control of the libraries, which were then sequenced on Illumina HiSeq^TM^ 2000. Raw reads produced by sequencing were stored in Fastq format. Raw reads with adaptors and unknown nucleotides and low quality reads with more than 20% low quality base calling (base quality ≤10) were removed to leave clean reads. Trinity (Haas et al., 2013) was employed for *de novo* assembly based on the clean reads to produce Unigenes.

To annotate the Unigenes, blast (blastX and blastn) searches were carried out against NR, Swiss-Prot, KEGG, COG and NT databases (e-value<0.00001). Annotations of the proteins with the highest similarity to each Unigene were retrieved. Then GO (Gene Ontology) annotations based on the similarity were assigned to each Unigene using Blast2GO (Conesa et al., 2005). Expression of Unigenes was calculated using the FPKM method (Fragments Per Kb per Million fragments) (Mortazavi et al., 2008). The formula is: FPKM=10^6^C/NL/10^3^, where C is the number of fragments that aligned specifically to a certain Unigene; N is the total number of fragments that aligned to all Unigenes; L is the base number in the CDS of the Unigene.

### Evaluation of genome quality

To evaluate the coverage of the assembly, all the paired-end Illumina short reads were mapped to the assembly using BWA (http://bio-bwa.sourceforge.net/) (Li and Durbin, 2009). Gene completeness was evaluated using the ESTs generated from RNA sequencing. The ESTs were mapped to the assembly using BLAT (http://genome.ucsc.edu/goldenpath/help/blatSpec.html). For CEGMA (Core Eukaryotic Genes Mapping Approach, http://korflab.ucdavis.edu/dataseda/cegma/) evaluation, we build a set of highly reliable conserved protein families that occur in a range of model eukaryotes (Parra et al., 2007). Then we mapped the 248 core eukaryotic genes to the genome. The genome was also assessed using BUSCO (Benchmarking Universal Single-Copy Orthologs: http://busco.ezlab.org/) gene set analysis (Simão et al., 2015), which includes 956 single-copy orthologous genes.

### Genome annotation

Repeat elements were annotated using a combined strategy. Alignment searches were undertaken against the RepBase database (http://www.girinst.org/repbase), then Repeatproteinmask searches (http://www.repeatmasker.org/) were used for prediction of homologs (Jurka et al., 2005). For *de novo* annotation of repeat elements, LTR_FINDER (http://tlife.fudan.edu.cn/ltr_finder/), Piler (http://www.drive5.com/piler/), RepeatScout (http://www.repeatmasker.org/) and RepeatModeler (http://www.repeatmasker.org/RepeatModeler.html) were used to construct a *de novo* library, then annotation was carried out with Repeatmasker (Price et al., 2005). Non-coding RNA was annotated using tRNAscan-SE (http://lowelab.ucsc.edu/tRNAscan-SE/) (for tRNA) or INFERNAL (http://infernal.janelia.org/) (for miRNA and snRNA). Since rRNA sequences are highly conserved among plants, rRNA from *S. baicalensis* was screened by blast searches.

Gene structure screening was carried out through a combination of homology, de novo and EST based predictions. A gene set including protein coding sequences from *Arabidopsis thaliana*, *Oryza brachyantha*, *Populus trichocarpa*, *Salvia miltiorrhiza*, *Sesamum indicum*, *Solanum lycopersicum* and *Utricularia gibba* was mapped to the assembly of *S. baicalensis* using blast (http://blast.ncbi.nlm.nih.gov/Blast.cgi) (E value≤1e^−5^) and gene structure of each hit was predicted through Genewise (http://www.ebi.ac.uk/∼birney/wise2/) (Birney et al., 2004). *De novo* gene structure identification was performed using Augustus (http://bioinf.uni-greifswald.de/augustus/) and GlimmerHMM (http://ccb.jhu.edu/software/glimmerhmm/). Gene structure was also screened by mapping ESTs to the assembly through Blat (http://genome.ucsc.edu/cgi-bin/hgBlat). All the resulting genes were corrected by reference to the transcriptome data and integrated into a non-redundant gene set using EVidenceModeler (EVM, http://evidencemodeler.sourceforge.net/) (Haas et al., 2008). The genes were further corrected with PASA (Program to Assemble Spliced Alignment, http://pasa.sourceforge.net/) to predict UTRs and alternative splicing (Haas et al., 2008), which generated 28,930 gene models.

Gene function was annotated by performing BLASTP (E-value ≤ 1e^−5^) against the protein databases, SwissProt (http://www.uniprot.org/), TrEMBL (http://www.uniprot.org/) and KEGG (http://www.genome.jp/kegg/). InterPro (https://www.ebi.ac.uk/interpro/) and Pfam (http://pfam.xfam.org/) were used for screening the functional domains of the proteins. Then GO (Gene Ontology) terms based on the searches were assigned to each gene.

### Data accessibility

Reference genome data are deposited in GenBank under project number PRJNA484052 and transcriptome sequence reads are deposited in the Sequence Read Archive (SRA) under accession number SRA accession SRP156996. The authors of the present manuscript reserve their priority for using these data for genome level analyses. By using these data, you agree not to use them for the publication of such genome level analyses without the written consent of the authors. If you agree to these terms, you can obtain the credentials for accessing the data by writing and/or email to Qing Zhao at Shanghai Chenshan Botanical Garden.

## Acknowledgements

This work was supported by National Natural Science Foundation of China (31870282, 31700268 and 31788103), Fund of Chinese Academy of Sciences (QYZDY-SSW-SMC026 and 153D31KYSB20160074), Chenshan Special Fund for Shanghai Landscaping Administration Bureau Program (G172402 and G162409) and by the CAS/JIC and Centre of Excellence for Plant and Microbial Sciences (CEPAMS) joint foundation for support to QZ, XYC and CM.

## Authour Contributions

QZ and CM initiated the programme, coordinated the project and wrote the manuscript. LJ, M-Y C, Y-M F, Y-K W and LY maintained and prepared the samples. QZ, W-Q Q, Z-C X and H-W S designed the sequencing strategy and performed sequencing. JY, QZ, X-Y C, Y-H H and CM performed analysis.

